# Warming increases richness and shapes assemblages of eukaryotic parasitic plankton

**DOI:** 10.1101/2024.12.31.630883

**Authors:** Amruta Rajarajan, Sławek Cerbin, Kingsly C. Beng, Michael T. Monaghan, Justyna Wolinska

## Abstract

**Background:** Anthropogenic activities have led to a global rise in water temperatures, prompting increased interest in how warming affects infectious disease ecology. While most studies have focused on individual host-parasite systems, there is a gap in understanding the impact of warming on multi-host, multi-parasite assemblages in natural ecosystems. To address this gap, we investigated freshwater eukaryotic parasite communities in ten natural lakes near Konin, Poland: five artificially heated and five non-heated “control” lakes. Since 1958, the heated lakes have experienced a mean annual temperature increase of 2 °C due to hot water discharge from two adjacent power plants. We collected seasonal environmental DNA (eDNA) samples from surface waters over a two-year period and applied targeted metabarcoding to compare the richness and distribution of eukaryotic parasites across lake types, with a focus on protists and fungi.

**Results:** Using literature searches and sequence metadata from GenBank, we identified putative parasites which included Alveolates, Stramenopiles, basal Fungi and Ichthyosporeans, as well as their associated hosts. Heated lakes harboured distinct parasite assemblages, with higher richness of chytrids and aphelids, suggesting thermal preferences among certain freshwater microeukaryotic parasites. Other groups exhibited clear seasonal trends, with richness of oomycetes peaking in spring and summer, and that of Cryptomycota in winter and autumn. A general linear model revealed a marginally positive correlation between chytrid parasite richness and richness of their green algal, diatom, and dinoflagellate hosts. Post-hoc analyses indicated that heated lakes exhibited greater seasonal variation in chytrid parasite richness and a stronger correlation between host and parasite richness than control lakes.

**Conclusion:** These findings demonstrate that warming can induce strong shifts in the richness and assemblages of freshwater microeukaryotic parasites. Using chytrids as a focal group, we additionally demonstrate that warming may amplify seasonal variation in parasite richness and strengthen host-parasite richness relationships.

## Introduction

Warming in aquatic environments has led to an average increase in freshwater temperatures of 0.34 °C per decade (O’Reilly et al., 2015). Elevated temperatures can drive deoxygenation (Jane et al., 2021), changes in seasonality and reduced ice cover (Grant et al., 2021), shifts in lake productivity dynamics (Gilarranz et al., 2022), and alterations in mixing regimes (Woolway & Merchant, 2019). These changes have cascading effects on species distributions and food webs (Bartley et al., 2019). The magnitude of warming-driven changes on aquatic life is unprecedented, highlighting a critical need to understand their broader ecological implications (IPCC 2023).

While the effects of warming on plankton have been widely studied due to their central role in nutrient cycling (Bakhtiyar et al., 2020; Naselli-Flores & Padisák, 2023), the parasites of plankton remain relatively understudied despite their significant ecological contributions. For instance, chytrid fungi (Chytridiomycota) infect various phytoplankton, including green algae (reviewed in Frenken et al., 2017), diatoms (reviewed in Danz & Quandt, 2023), and dinoflagellates (Fernández-Valero et al., 2023, 2024). By producing motile zoospores, chytrids create additional trophic links (mycoloops) that sustain zooplankton populations (Abonyi et al., 2024; Kagami et al., 2014). Chytrid outbreaks can exert top-down control on toxic cyanobacterial (Gleason et al., 2015) and dinoflagellate blooms (Lepelletier et al., 2014). Other parasitic groups, such as Cryptomycota, hyperparasitize chytrids (James et al., 2013; Powell & Letcher, 2019), while Aphelidiomycota (aphelids) (Jephcott et al., 2017) and Oomycota (oomycetes) (Thines, 2018) infect various phytoplankton hosts. Perkinsea, a group previously considered to exclusively contain intracellular parasites of green algae, was recently discovered to be more diverse, with potential roles in ecosystem processes beyond parasitism (Metz et al., 2023). Other highly diverse parasite groups include Apicomplexans (Gad et al., 2023) and Cercozoans (Lefèvre et al., 2008), although their contribution to aquatic ecosystem processes is less understood.

The influence of warming on infectious disease extends beyond parasite development to affect host abundance, physiology, and immunity (King et al., 2023). For instance, a four-month indoor mesocosm study found that 4 °C elevation in spring temperature (to 21 °C) shortened the bloom duration of the diatom *Synedra* sp., resulting in an earlier end to the epidemic of its chytrid parasite *Zygorhizidium planktonicum* (Frenken et al., 2016). Conversely, an outdoor mesocosm study of amphipod host communities exposed to the generalist trematode parasite *Maritrema novaezealandensis* under a 4 °C summer temperature increase (to 25 °C) showed elevated infection prevalence, along with reduced host species richness and abundance (Mouritsen et al., 2018). These findings demonstrate how warming can impact disease dynamics in individual host-parasite systems within aquatic environments.

Parasite species richness is integral to ecosystem health (Hudson et al., 2006) is primarily influenced by the host distribution and richness of their hosts in distinct ways. Chytrid parasite distributions are closely linked to the spatial co-occurrence of their green algal and diatom hosts (Ilicic et al., 2024). More broadly, parasite richness also correlates positively with host richness across biomes, according the host-diversity-begets-parasite-diversity hypothesis (Kamiya et al., 2014; Krasnov & Poulin, 2015). For example, ponds with higher amphibian host richness harbored higher richness of trematode parasites (Johnson et al., 2016). Changes in host and parasite species richness can, in turn, affect disease dynamics in complex ways. In one case, elevated amphibian host richness reduced infection prevalence of the trematode *Ribeiroia ondatrae* (dilution effect) but amplified parasite competition (Johnson et al., 2013).

Despite these insights, our understanding of how global warming affects parasite richness remains incomplete (Sures et al., 2023). Warming-driven changes in species assemblages can lead to ‘ecological fitting’ due to novel host-parasite encounters and the resulting increased likelihood of host-switching (Wells & Clark, 2019). This could potentially increase phenotypic and genotypic diversity of parasites (Agosta et al., 2010). Moreover, anthropogenic influence can have unexpected impacts on the generally positive correlation between host and parasite richness in natural ecosystems. For instance, islands that experienced extensive fishing exhibited higher fish host richness, but lower helminth parasite richness, compared to unfished islands (Wood et al., 2018). In contrast, warming-driven habitat loss of host species reduced helminth parasite richness (Carlson et al., 2017). These inconsistencies highlight the need for long-term studies of multi-host, multi-parasite systems to better understand the role of warming in shaping disease ecology.

One promising approach to study the effects of warming on parasite assemblages involves *in situ* sampling of natural lakes. The Konin lake complex in Poland offers an ideal opportunity. Since 1958, five of these lakes have been artificially warmed by hot water discharge from two nearby power plants, resulting in altered abiotic conditions and changes in plankton community composition compared to the five nearby non-heated (hereafter “control”) lakes (Zdanowski et al., 2020). The heated lakes are on average 2 °C warmer (Beng et al., 2023), mimicking the projected global freshwater temperature increases by 2100 (IPCC 2023). Additionally, these lakes experience reduced winter ice cover (Socha & Zdanowski, 2001). Warming has led to shifts in *Daphnia* species composition in these lakes, favoring the more heat-tolerant *D. galeata* (Dziuba et al., 2020). Moreover, the heated lakes exhibit higher richness and distinct community composition of various phytoplankton (green algae, protists, fungi) and zooplankton (crustaceans, rotifers) taxa (Beng et al., 2023; Ejsmont-Karabin et al., 2020). There is also an increased abundance of invasive species, including the tape grass *Vallisneria spiralis* (Socha & Hutorowicz, 2009) and the mussels *Corbicula fluminea* (Urbańska et al., 2018) and *Sinanodonta woodiana* (Urbańska et al., 2019). *Sinanodonta woodiana* occurrence further favors the presence of its parasitic worm *Aspidogaster conchicola* (Yuryshynets & Krasutska, 2009). Heated lakes also harbor introduced species of fish (Kapusta & Bogacka-Kapusta, 2015). These changes in environmental conditions and host community composition provide a valuable opportunity to investigate the effects of warming on parasite diversity in natural settings. Although warming reduces *Caullerya mesnili* prevalence in *Daphnia* hosts (Dziuba et al., 2023), comprehensive investigations into differences in parasite assemblages between heated and control lakes are still lacking.

This study leverages this unique set of artificially heated and control lakes in Poland to explore ca. 60 years of warming effects on eukaryotic parasitic plankton. Using eDNA metabarcoding of seasonally collected surface water samples (winter, spring, summer, autumn) over two years from five heated and five control lakes to address the following questions: (i) Which parasites are present in these lakes? (ii) Do certain parasites exhibit a preference for habitat types (heated or non-heated), leading to distinct assemblages in heated versus control lakes? (iii) Does parasite richness differ between heated and control lakes? and (iv) Does parasite richness correlate with host richness?

## Material and methods

### Study sites and eDNA sampling

Information on study sites and the timing and locations of eDNA sampling is adapted from Beng et al., (2023). This study includes five thermally altered (“heated”) lakes (Gosławskie, Licheńskie, Mikorzyńskie, Pątnowskie and Ślesińskie) and five control lakes (Budzisławskie, Gopło, Skulskie, Skulska Wieś and Suszewskie). Each lake was sampled once per season throughout 2020 and 2021, with the exception that no summer samples were obtained in 2021 and all lakes were sampled twice in autumn 2020 (Fig. S1, Table S1). All lakes, except for the control lakes Suszewskie and Budzisławskie, are connected by water channels, potentially enabling dispersal, migration and colonization (Dziuba et al., 2017; Hillbricht-Ilkowska & Zdanowski, 1988).

Water from the lakes Gosławskie, Licheńskie, Mikorzyńskie, Pątnowskie and Ślesińskie has been used to cool lignite-combusting power plants situated in Konin (since 1958) and Pątnów (since 1970), with the heated effluent subsequently discharged back into the lakes. This discharge occurs year-round in all heated lakes except Ślesińskie, which receives thermal effluent only from May to September. On average, heated lakes exhibit water temperatures 2 °C higher than control lakes. Beyond temperature, these lakes also show elevated concentrations of total and soluble reactive phosphorus, particularly in the autumn (Beng et al., 2023). While used for recreation and fish-farming, regular monitoring confirms that hot water discharge does not cause chemical pollution. All study lakes are subject to some level of anthropogenic disturbance, including agricultural runoff, aquaculture and urbanization.

To obtain eDNA samples, 2 L of surface water was collected at each sampling event using a clean HDPE bottle. Water was filtered through a glass fiber mesh (Whatman GF/F, 25 mm diameter, 0.7 µm pore size) using vacuum filtration at 200 mbar. Filters were first frozen at - 80 °C, freeze-dried at -45 °C for 8 hours and finally stored at -20 °C until DNA extraction.

### Generation of sequence data

#### Molecular laboratory procedures and sequencing

Molecular laboratory procedures are reproduced from Beng et al., (2023). Filters containing eDNA were pulverized by adding one stainless steel bead (5 mm diameter, Qiagen GmbH, Hilden, Germany) and tissueLyser (Qiagen), and shaking three times for 90 seconds at 30 Hz in a mixer mill MM301 (Retsch GmbH, Haan, Germany). The tubes were briefly centrigfuged to collect the material into a pellet. DNA was extracted using the NucleoSpin Plant II extraction kit (Sinhahery-Nagel, Düren, Germany), following the manufacturer’s protocol, and stored in TE buffer at -20 °C until further processing.

For DNA amplification, two primer sets were used: “EUK15” TAReuk454FWD1: 5’-CCAGCASCYGCGGTAATTCC-3’; TAReukREV3: 5’-ACTTTCGTTCTTGATYRA-3’; both primers from Stoeck et al., (2010), targeting a fragment of the 18S rRNA gene of eukaryotes, and “CHY” ITS4ngsF: 5’-GCATATCAATAAGCGSAGGA-3’ (Van den Wyngaert et al., 2022); LF402R: 5’-TTCMCTTTNMRCAATTTCAC-3’ (Thomé et al., 2024), targeting a fragment of the 28S rRNA gene of fungi. For the “EUK15” primer set, 10 ng template DNA was used in 25-µL PCR reactions with the following conditions: an initial denaturation at 95 °C for 30 seconds, followed by 30 cycles (95 °C for 30 s, 45 °C at 30 s, 72 °C at 30 s), with a final extension of 72 °C for 5 minutes. PCR products were normalized to 5 ng/µL and used as templates for the second PCR, where Illumina Nextera indexes were added to 25-µL PCRs under these conditions: 95 °C for 2 minutes, followed by 8 cycles (95 °C for 20 s, 52 °C for 30 s, 72 °C for 30 s), and a final extension at 72 °C for 3 minutes. For the “CHY” primer set, 10 ng template DNA was used in 25-µL PCRs with the following conditions: an initial denaturation at 96 °C for 30 seconds, followed by 20 cycles (90 °C for 30 s, 50 °C for 30 s, 72 °C for 60 s), with a final extension of 72 °C for 3 minutes. 1 µL of PCR product was used as template to repeat the enrichment PCR using the same conditions except that the final extension at 72 °C was for 5 minutes. PCR products were purified using a magnetic bead protocol (Agencourt AMPure XP, Beckman Coulter, Indianapolis, IN, USA), following the manufacturer’s instructions. DNA was quantified using a Quantus fluorometer and QuantiFluor dsDNA system (Promega, Madison, WI, USA) and PCR products were normalized to a concentration of 5 ng/µL for use in indexing. Indexing PCR was performed with the following cycling conditions: initial denaturation at 95 °C for 2 min, followed by 8 cycles (95 °C for 20 s, 52 °C for 30 s, 72 °C for 30 s), with a final extension at 72 °C for 3 min. Barcoded samples were pooled in equimolar concentrations and paired-end sequenced on the Illumina MiSeq (600 cycles, v3 chemistry) platform at the Berlin Center for Genomics in Biodiversity Research (BeGenDiv). Each sequencing run included one no-template negative control.

#### Bioinformatics

Each library (EUK15 and CHY) was sequenced in two runs: the first run included samples from winter 2020, spring 2020, summer 2020 and autumn 2020, while the second run included samples from winter 2021, spring 2021, autumn 2021 and an additional set of samples taken in autumn 2020. Bioinformatic processing was performed as described in Beng et al., (2023). The first sequencing run yielded 2.4 million raw reads for EUK15 and 1.9 million raw reads for CHY (Beng et al., 2023), while the second run yielded 643,742 raw reads for EUK15 and 741,018 raw reads for CHY. Primer sequences were removed with cutadapt (Martin, 2011) and reads were processed using the package DADA2 v 1.28 (Callahan et al., 2016) in R v 4.3.0 (R Core Team, 2023). Forward and reverse reads were respectively: filtered with maximum expected errors set to 2.6 and 2, then truncated at 260 bp and 200 bp respectively before merging (see separate supplementary R scripts). Quality filtering and *de novo* chimera removal retained 59% of EUK15 and 58.3% of CHY reads. Reads from the two runs were combined into a “protist dataset” (EUK15) and a “fungal dataset” (CHY). Amplicon Sequence Variants (ASVs) were identified for each dataset using the *dada* function, and ASV-count tables were prepared. Taxonomic assignment of ASVs in the protist dataset was conducted using the Protist Ribosomal database PR2 v.5.0.0 (Guillou et al., 2013). For the fungal dataset, taxonomic assignment was performed using BLAST+ with a custom database (see below).

Despite the lower sequencing depth in the second runs, all rarefaction curves plateaued (Fig. S2), confirming sufficient sequencing depth. Thus, all samples were retained for further analyses without rarefaction. Contaminant ASVs, defined as those present in negative controls, were removed using the phyloseq package in R (McMurdie & Holmes, 2013). Specifically, 17 contaminant ASVs were removed from the first run of the protist dataset, while 3 and 2 contaminant ASVs were removed from the first and second runs of the fungal dataset, respectively, comprising <0.01% reads in each dataset. Bioinformatic processing resulted in 3297 ASVs in the protist dataset and 4243 ASVs in the fungal dataset.

### Classification of ASVs as parasites

ASVs were classified as parasites using a modified approach from Beng et al. (2021) (detailed in Fig S3). Two custom parasite sequence databases were created for this purpose: one containing sequences from the NCBI nucleotide database representing 10 higher taxa discovered in the protist dataset (Fig. S4), and the other containing NCBI sequences of 9 taxa discovered in the fungal dataset (Fig. S5). Additionally, we incorporated (i) ribosomal sequences from NCBI (18S, 28S, and ITS), (ii) sequences from parasite groups reported to occur in the Konin lake complex (Cestoda, Nematoda, Monogenea, Digenea, Trematoda and Acanthocephala), which infect fish (Pojmańska et al., 2012), mussels (Yuryshynets & Krasutska, 2009), and snails (Cichy et al., 2020), and (iii) sequences of eukaryotic taxonomic groups known to contain parasite species, identified through Web of Science literature searches performed for each unique phylum listed in the Fungal Names database (F. Wang et al., 2023), using the terms “[Phylum name] AND [parasit* OR pathogen*]” (Table S2). Duplicate sequences were removed using seqkit (Shen et al., 2016). Each ASV was then identified to species- or higher taxonomic level using its respective custom database and BLAST+ on a high-performance computing cluster (Bennett et al., 2020). Only the top ten hits with ≥90% sequence similarity and 100% query cover were processed further. For each ASV, “host” entries within GenBank metadata of the top-bitscore BLAST hit (Beng et al., 2021) were extracted using the rentrez package (Winter, 2017).

Species names from the protist and fungal datasets (for ASVs that were assigned to species level) were used for literature searches on Web of Science with the search terms “[Species name] AND [parasit* OR pathogen* OR infect*]”. Relevant articles were checked for evidence of parasitism and host association. BLAST hits of ASVs with no associated host entries in GenBank or host information from literature searches were classified as non-parasitic (Fig. S3).

Host entries obtained from GenBank were manually categorized into host groups (see Fig. 1). Only ASVs associated with live hosts were classified as parasites; those associated with abiotic environmental samples (e.g., water, soil, mud sediment) or non-living substrates (e.g., “decaying log”, “fallen branch”) were excluded. ASVs representing symbionts (e.g., algal endosymbionts of protists, lichen-associated ASVs, dinoflagellate symbionts of sponges) or organisms within the pathobiome of diseased hosts were also excluded. This rigorous classification ensured that only ASVs with confirmed parasitic lifestyles were included in the final analysis, enabling an accurate assessment of parasite richness and distribution across the studied lakes (Fig. S3).

**Figure 1.**
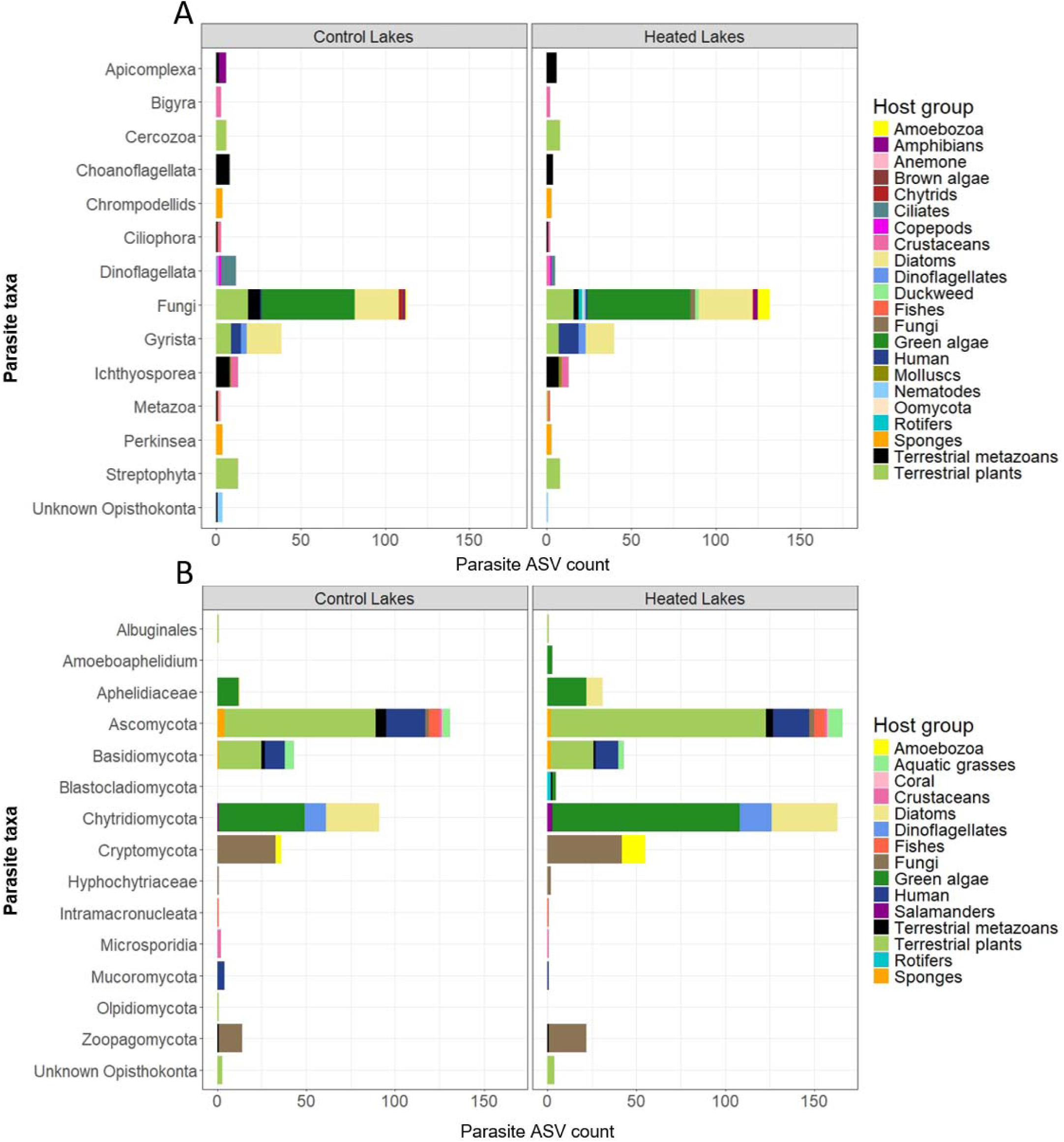
Parasite ASV count in the (A) protist and (B) fungal datasets. Colors represent broad host taxa to which identified hosts of the parasite ASVs belong (see Methods and Fig. S3 for details). Note that Fungal ASVs from the protist dataset were not considered for statistical analyses.

### Statistical analyses

Individual lakes of each type (heated, control) were considered replicates in all statistical analyses, but included independent sampling events over two consecutive years (Table S1). Sample sizes differed for the seasons in our analyses: winter (n = 10), spring (n = 10), summer (n = 5) and autumn (n = 15).

#### ASVs indicative of heated or control lakes

To assess whether certain parasite ASVs were indicative of either heated or control lakes, we used the *signassoc* function (9999 permutations) from the indicspecies package in R. This analysis was performed using presence-absence data for parasite ASVs (De Caceres & Legendre, 2009) (mode=0). Multiple testing correction was implemented using the Sidak method within the indicspecies package.

#### ASV richness differences in dominant parasite groups between heated and control lakes

Parasite groups were scrutinized to identify those associated with aquatic hosts (see Fig. 1). The four parasite groups with the highest ASV count were selected for comparisons across lake types and seasons. For example, in the protist dataset, 66 ASVs were identified as Gyrista (Stramenopiles phylum), but only 31 Gyrista ASVs belonged to Peronosporomycetes, after excluding terrestrial hosts. Consequently, only Peronosporomycetes (=Oomycetes, Subdivision: Gyrista), representing 7.9% of ASVs in the protist parasite dataset, were included in ASV richness analyses. Additionally, 216 ASVs in the protist dataset were identified as fungi but only fungi from the fungal dataset were considered for further analyses. In the fungal dataset, 297 and 86 ASVs belonged to Ascomycota and Basidiomycota, respectively, but 86.8% and 87.2% of these ASVs were associated with terrestrial hosts (such as metazoans, plants and humans; see Fig. 1B) and were therefore excluded. Chytrids (Phylum: Chytridiomycota), aphelids (Phylum: Aphelidiomycota) and Cryptomycota infecting aquatic hosts were analyzed, collectively representing 50.3% of ASVs in the fungal parasite dataset.

The ASV richness for each selected group was log-transformed and analyzed using a Generalized Linear Mixed Model (GLMM). The most parsimonious model with the best fit was used to examine parasite richness across lake types, seasons and years: parasite ASV richness ∼ lake type * season + (1 | lake) + (1 | sequencing run) + (1 | year), family = poisson, using the *glmer* function of the lme4 package (Bates et al., 2024) in R. The significance of fixed effects was assessed with a Type III ANOVA using the *Anova* function of the car package (Fox et al., 2019). For chytrid parasite richness, a separate model was employed to investigate its correlation with host richness (see below).

#### Correlation of chytrid parasite richness and host richness

A second GLMM was used to test whether differences in chytrid parasite richness correlated with differences in host richness across heated and control lakes (Beng et al., 2023). Here, “chytrid parasite richness” was defined as the count of chytrid parasite ASVs in the fungal dataset, while “host richness” referred to the count of ASVs from the Subdivisions Chlorophyta, Charophyta, Chromista, Gyrista, and Dinoflagellata in the protist dataset. These host groups were selected because 98% of chytrid parasite ASVs were associated with one or more of these groups (see Fig. 1B). 2% chytrid parasite ASVs were excluded from this analysis because they associated with salamander hosts, and the richness of salamander hosts was not surveyed (see Table S3). Both host and parasite richness values were log-transformed. We then used the following GLMM structure: chytrid parasite richness ∼ lake type * season + host ASV richness + (1 | lake) + (1 | sequencing run) + (1 | year), family = poisson. Fixed effects were evaluated with a Type III ANOVA. After observing significant effects of host richness, lake type, season and lake type × season interaction on parasite richness (see Results), we calculated confidence intervals and visualized a general linear model for host and parasite richness, separately for each lake type (see Fig. 4). The model was plotted using the “geom_smooth” argument (method=”glm”, se=TRUE) from the ggplot2 package (Wickham, 2016) in R.

After discovering the significant effect of lake type, host richness and the lake type × season interaction on chytrid parasite richness (see Table 1), we computed partial R^2^ values to assess the effect sizes of host richness and season on parasite richness. This was done separately for heated and control lakes. Random effects were excluded from this test because mixed-effects structures resulted in a singular fit. Instead, GLMs were specified for each lake type with the structure: parasite richness ∼ host richness + season. Partial R^2^ values for each fixed effect were computed using the rsq package in R (Zhang, 2024).

**Table 1.**
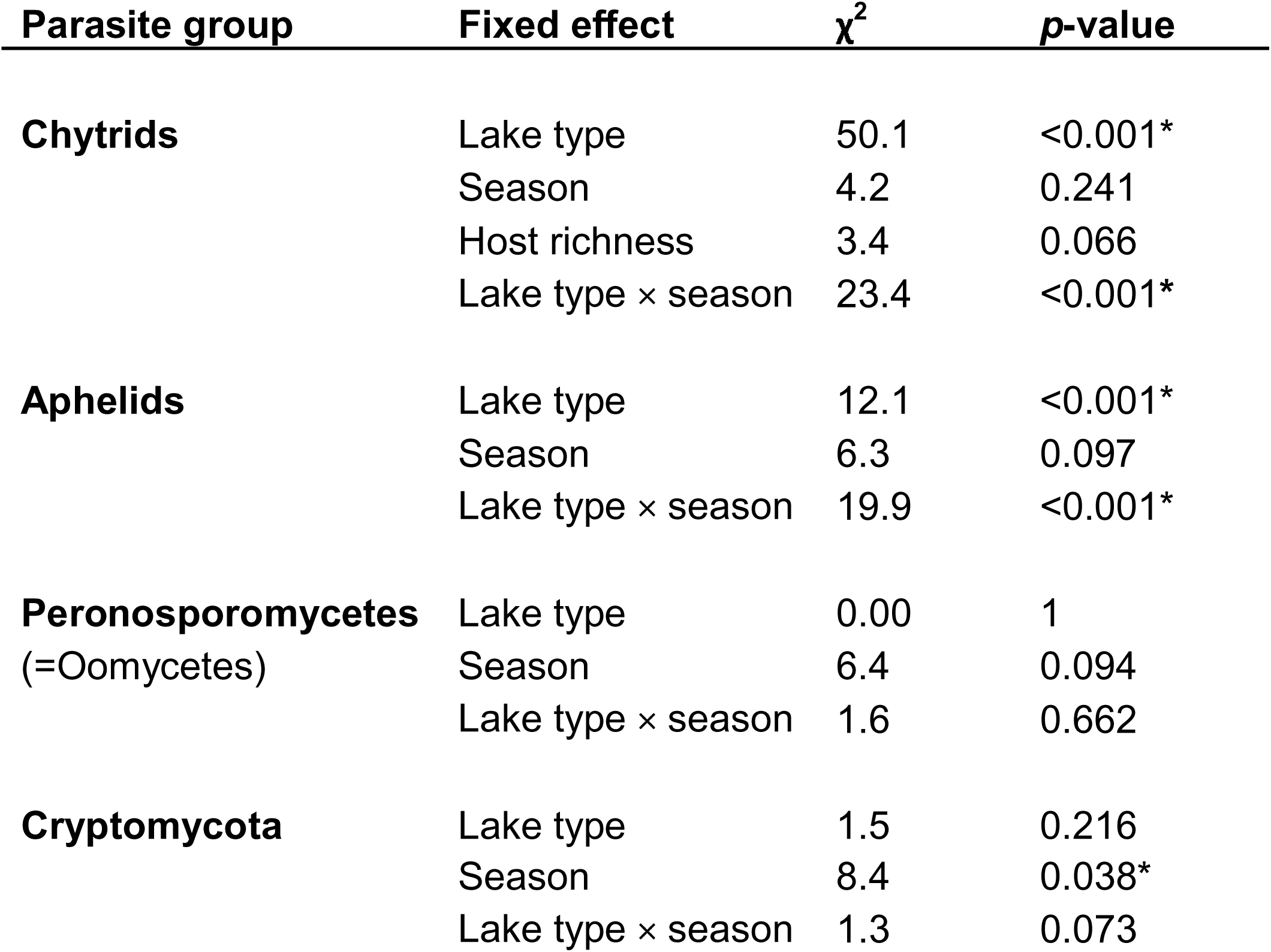
Results of a type III ANOVA of GLMM investigating ASV richness for the four most ASV-diverse parasite groups across lake type and season. Lake and sequencing run were specified as random effects. The chytrid parasite richness model uniquely incorporated host richness (Chlorophyta, Charophyta, Chromista, Gyrista, and Dinoflagellata) as a predictor. Significant results (\**p*<0.05) are marked with an asterisk.

## Results

### Characterization of parasite assemblages and hosts in the protist and fungal datasets

In the protist dataset, 390 ASVs (12% of the dataset) were classified as parasites. These included: 43 ASVs confirmed as parasitic based on both literature evidence and GenBank host entries, 55 ASVs classified as parasites using literature evidence alone, and 292 ASVs classified based solely on GenBank host entries (Table S2). A total of 151 parasite ASVs (39%) were identified to the species level, representing 54 species from 23 genera (Table S3), which were further grouped into 14 higher taxa (Fig. 1A). The highest ASV richness was observed among Fungi, Gyrista, and Ichthyosporea (Fig 1A). Most parasite ASVs (66%) were associated with hosts such as green algal, diatom, or terrestrial-plant hosts (Fig. 1A, Table S3).

In the fungal dataset, 683 ASVs (16% of the dataset) were identified as parasites: 198 ASVs were classified as parasitic using both literature evidence and GenBank host entries, 298 ASVs based on GenBank host entries alone, and 187 ASVs using literature searches of species names. A total of 496 ASVs (73% of parasite ASVs) showed >90% sequence similarity and 100% query cover with known species in GenBank (Table S4). Based on these criteria, the fungal dataset contained 268 putative species across 218 genera, classified into 15 higher taxa. The highest ASV richness was observed among Ascomycota, Chytridiomycota and Cryptomycota (Fig. 1B). Similar to the protist dataset, a notable proportion of parasite ASVs (45%) in the fungal dataset were linked to hosts such as green algae, diatoms, or terrestrial plants (Fig. 1B, Table S4).

### ASVs indicative of lake type

A total of 39 parasite ASVs were indicators of lake type: 5 ASVs were specific to control lakes, and 34 ASVs were specific to heated lakes (Fig. 2). Among these were 19 chytrids, 8 aphelids, 4 Cryptomycota, 7 protists from diverse lineages, and 1 metazoan (Table S5). Control lakes indicators included chytrids ASVs (263 and 34) from unclassified members of the Order Rhizophydiales. In contrast, heated-lake indicators comprised various taxa, including chytrid species *Dinomyces arenysensis*, *Dangeardia mamillata*, *Rhizophydium chlorogonii*, *Quaeritorhiza haematococci* and an unknown Rhizophydiales parasite (ASV 210). Other heated-lake indicators were *Elliptio complanata* (Unionidae mussel), *Ustilago esculenta* (Basidiomycete fungus) along with unclassified ASVs from Dinophyceae (ASV 110) and Peronosporales fungi (ASV 286). Additionally, *Aphelidium desmodesmi* and four unclassified ASVs from the Family Aphelidiaceae (ASVs 60, 220, 194 and 101) showed a preference for heated lakes.

**Figure 2.**
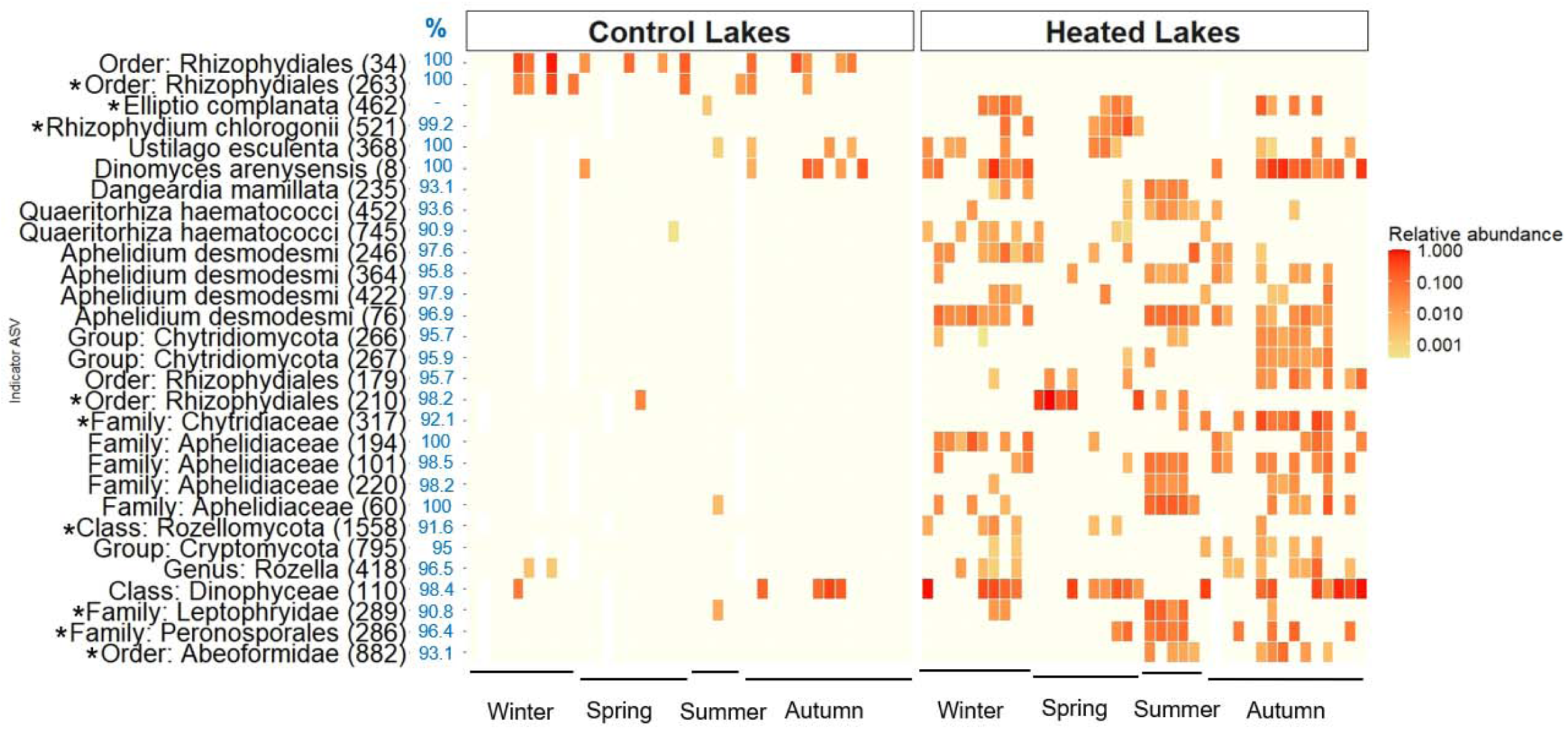
Relative abundances of ASVs indicative of control (left panel) and heated (right panel) lakes. Each square represents the relative abundance of indicator ASVs (rows) in individual lakes (columns), grouped by the season of sampling indicated on the x-axis. Sampling occurred over two consecu ive years (See Methods). Of the 39 indicator ASVs identified (Table S5), 29 are presented here, as they were present in more than 20% of samples. ASV numbers are specified in brackets. For ASVs in the protist dataset (marked with an asterisk), the lowest taxonomic classification assigned using the PR2 database is reported, with values in the blue “%” column indicating sequence similarity to the BLAST hit used to obtain host information (host entry from GenBank metadata). For ASVs from the fungal dataset, the lowest known classification from the BLAST hit is reported, with the blue “%” values indicating sequence similarity between the ASV and the BLAST hit. The classification of *E. complanata* as a parasite was based on a literature search (Table S5).

### Parasite ASV richness

Among the four parasite groups with the highest ASV richness - chytrids, aphelids, Peronosporomycetes (Oomycetes), and Cryptomycota - chytrids and aphelids had significantly higher ASV richness in heated lakes (Fig. 3, Table 1), with a significant interaction between lake type and season. Chytrid ASV richness peaked in heated lakes during autumn and summer (Fig. 3A). Aphelid richness was higher in heated lakes in all seasons except spring (Fig. 3B). Peronosporomycete ASV richness varied seasonally, with higher values in spring and a peak in summer (Fig. 3C). Both Peronosporomycetes and Cryptomycota exhibited lower ASV richness in 2021 compared to 2020 (Fig. 3C, D, Table 1).

**Figure 3.**
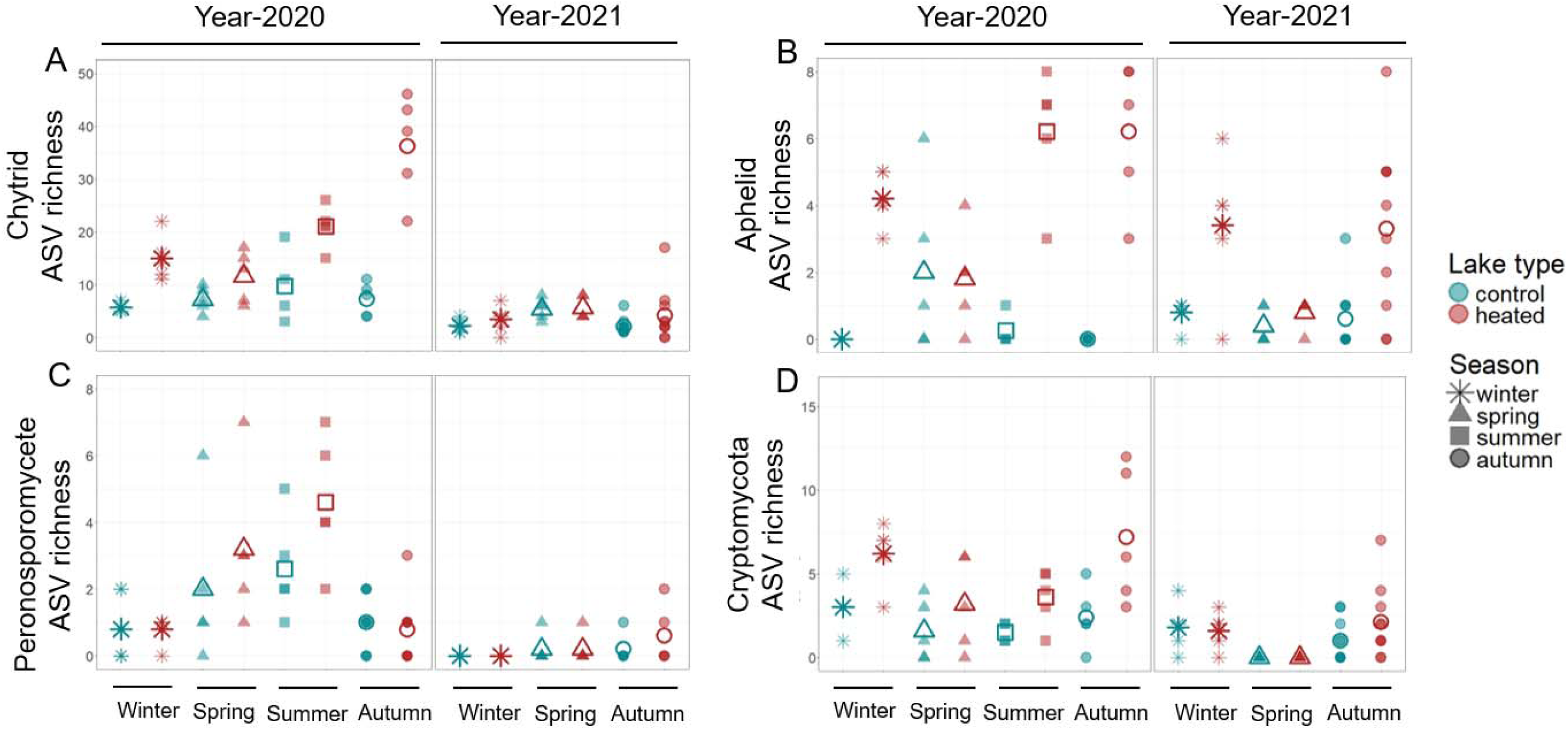
ASV richness of the four most ASV-rich parasite groups: (A) chytrids, (B) aphelids, (C) Peronosporomycetes (=Oomycetes) and (D) Cryptomycota. Colors indicate lake type and symbols indicate the season of sample collection. Filled symbols denote individual data points, whereas larger open symbols indicate seasonal mean values. Note that the y-axis scales differ across panels to optimize visualization.

### Correlation between host and parasite richness

Chytrid parasite richness showed a marginally significant positive correlation with host richness (Table 1, Fig. 4). Heated lakes harbored particularly high host richness during summer (Fig. 4), which aligned with the observed elevated chytrid richness for that season (Fig. 3A). However, when examining the correlation between parasite and host richness separately within each lake type, a deviation from the linear model was found: chytrid richness was exceptionally high in heated lakes, despite comparable host richness between heated and control lakes (Fig. 4). This discrepancy aligned with the significant interaction between lake type and season.

**Figure 4.**
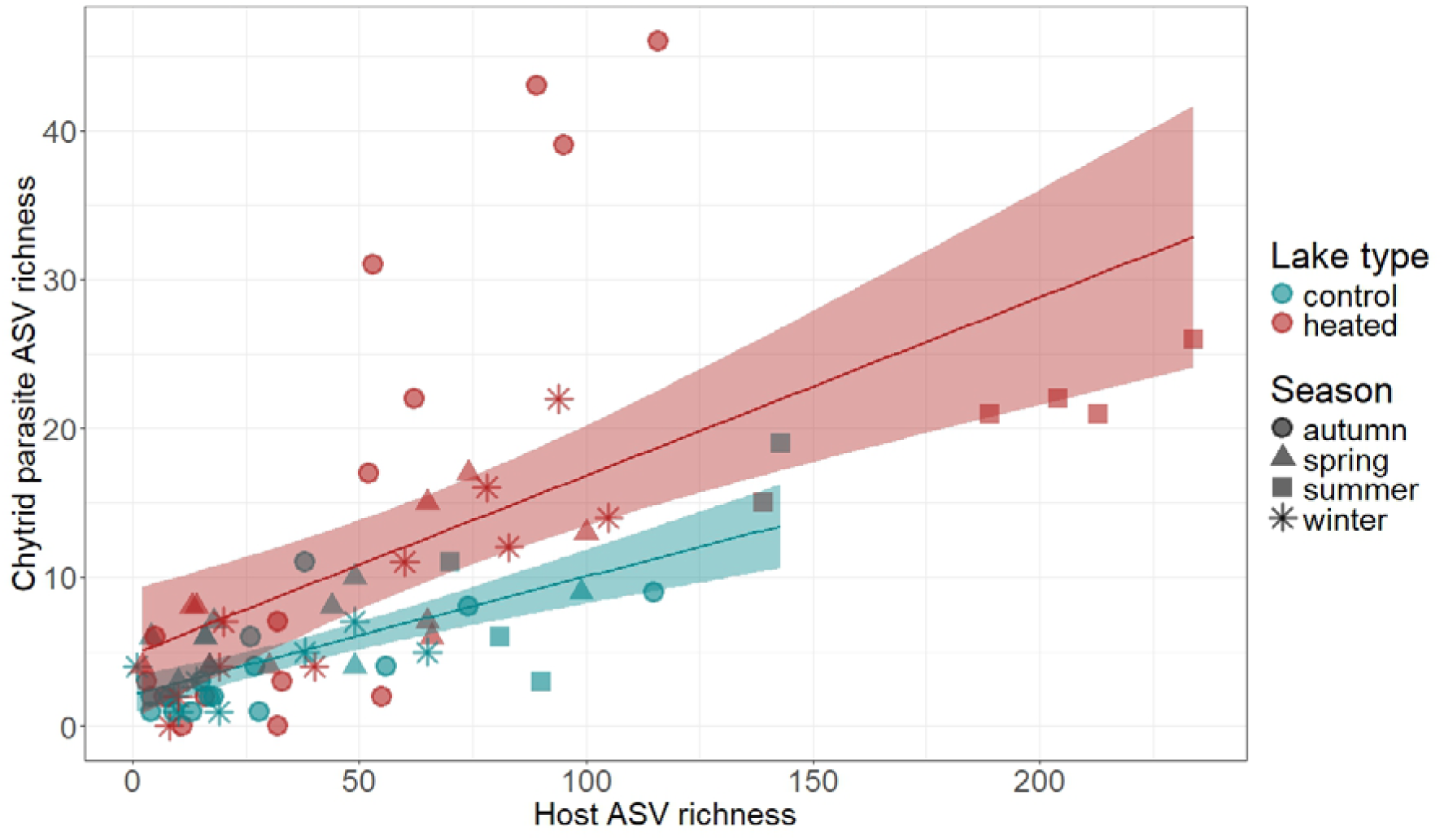
Correlation between chytrid parasite ASV richness (from the fungal dataset) and host ASV richness across Chlorophyta, Charophyta, Chromista, Gyrista, and Dinoflagellata (from the protist dataset). Shaded areas represent 95% confidence intervals. Colors differentiate between lake types, and symbols indicate the season of sample collection.

Post-hoc tests conducted separately for each lake type showed that in heated lakes, both host richness (partial R^2^= 0.72, *p* < 0.05) and season (partial R^2^= 0.58, *p* < 0.05) strongly correlated with variation in parasite richness. In control lakes, the correlation between parasite richness and host richness (partial R^2^= 0.32, *p* < 0.05) and season (partial R^2^= 0.11, *p* < 0.05) was less pronounced (Table S6). These results indicated that while host richness and season significantly impacted chytrid parasite richness in both lake types (Table S6), heated lakes exhibited stronger seasonal variation and a higher correlation with host richness.

## Discussion

Heated lakes received hot water discharge since 1958 and are on average 2 °C warmer, allowing us to investigate the long-term effects of warming on parasite assemblages. We identified parasites from seasonally sampled eDNA to assess differences in parasite species richness and assemblages between five artificially heated and five non-heated control lakes over two years, using parasite-specific sequence databases, their metadata, and literature. Our rigorous approach of classifying ASVs as parasites identified a diverse array of confirmed aquatic parasites, spanning major eukaryotic taxa Alveolates (Apicomplexa, Ciliophora, Dinoflagellata), Stramenopiles (Bigyra, Oomycetes), basal fungi (Chytrids, Rozellomycota) and Ichthyosporeans, highlighting the undocumented parasite diversity in natural lakes.

We found that certain parasite species exhibit a strong preference for either heated or control habitats, indicating that long-term warming can shape assemblages of parasite species. Chytrid and aphelid parasite richness was notably higher in heated lakes, supporting the idea that warming can enhance genotypic richness of parasites (Agosta et al., 2010). However, this trend was not observed for Peronosporomycetes (=Oomycetes) or Cryptomycota, which instead showed similar seasonal patterns of richness in all lakes (Peronosporomycete richness peaked in spring and summer, Cryptomycota richness peaked in autumn and winter). As predicted by the host-diversity-begets-parasite-diversity hypothesis (Kamiya et al., 2014; Krasnov & Poulin, 2015), chytrid parasite richness showed a positive correlation with host richness, but warming altered the strength of the correlation. Post-hoc analyses revealed that season and host richness correlated more strongly with parasite richness in heated lakes than control lakes, suggesting that warming may both amplify seasonal variation and increase the strength of correlation between host and parasite richness. Overall, these findings support the idea that thermal conditions strongly influence the composition and dynamics of parasite communities, possibly through a combination of direct effects on parasites and indirect effects via hosts.

### The Konin lake complex harbors ecologically relevant parasites, though a majority are unidentified

Our study identified 269 parasites that were assigned to species level in the protist dataset, or were >98% similar to a known parasite in GenBank. However, 86% of parasite ASVs in the protist dataset and 53% parasite ASVs in the fungal dataset could not be identified to species level (see Tables S3 & S4). This highlights the acute shortage of taxonomic information and reference sequences for freshwater parasites. Despite limited research, microeukaryotic parasites are known to impact ecosystem function. For example, zoospores of the chytrid *Zygorhizydium affluens* released from parasitized diatom blooms, provide a nutritious food source for *Daphnia*, thereby establishing a trophic link between producers and primary consumers (Kagami et al., 2007). In addition to *Z. affluens* (ASVs 31 and 717 in the current study), other parasite ASVs were >95% similar to known but as yet uncharacterized chytrid parasites, such as the chitin-degrading *Rhizoclosmatium pessaminum* (ASV 1901) (Powell et al., 2019), *Endochytrium ramosum* (ASVs 1418 and 2290), and *Staurastromyces oculus* (ASVs 1236, 1328 and 3041). These chytrids parasitize the diatom *Asterionella* (Rad-Menéndez et al., 2018), and green algae *Chara*, *Cladophora* (Simmons et al., 2020), and *Straurastrum* (Van den Wyngaert et al., 2017, see Table S43). Other ecologically relevant parasites detected in our study included two ASVs (181 and 1195; see Table S3) that showed >98% sequence similarity with *Psorospermium haeckeli* (Class: Ichythyophonida), a parasite of the crayfish *Ascatus ascatus* (Rug & Vogt, 1994), a host species that is impacted by the devastating Europe-wide crayfish plague (Wiśniewski et al., 2020). We also identified four ASVs (1156, 2064, 1231, and 1305, see Table S3) of the Apicomplexan parasite *Nematopsis temporariae*, which infects a range of amphibian hosts (Chambouvet et al., 2016).

ASV 286 (Family: Peronosporales) showed a strong preference for heated lakes (see Fig. 2) and exhibited 96.4% sequence similarity with *Pythium insidiosum*, a generalist oomycete parasite known to cause devastating root rot in commercial crops (Golińska & Świecimska, 2020), as well as disease in mammals (Mendoza, 2009) including humans (Mendoza et al., 1993). *Pythium* releases motile zoospores from infected plant roots or skin lesions (Mendoza et al., 1993). The close similarity of ASV 286 to a clinical sample from a patient with pythiosis (see Krajaejun et al., (2014) and GenBank BBXB02000196) suggests that heated lakes might function as a reservoir, providing suitable habitats or alternative hosts for this parasite.

We additionally found three parasite ASVs that showed 90-93% sequence similarity to *Batrachochytrium salamandrivorans* (*Bsal*), of which ASV 764 was predominantly detected in winter within heated lakes Gosławskie, Licheńskie, Mikorzyńskie, and Pątnowskie (see Table S5, Fig. S6). *Bsal* is responsible for global chytridiomycosis outbreaks in amphibians (Balàž et al., 2017; Laking et al., 2017; Yap et al., 2015). *Bsal* produces motile, water-borne transmission stages (zoospores) that are active within a thermal range of 10-15 °C but capable of surviving at temperatures as low as 5 °C (Martel et al., 2013). Consequently, epidemics are generally observed during colder months, such as late autumn, winter or spring (Schulz et al., 2020). Geospatial models predict a reduction in climatically suitable habitats for *Bsal* under global warming scenarios (Grisnik et al., 2023). However, recent chytridiomycosis outbreaks at water temperatures as high as 26.4 °C (Laking et al., 2017) suggest a broader thermal niche than previously anticipated. Although *Bsal* infections have so far not been detected in Poland, its sister species *B. dendrobatidis* is widespread (Palomar et al., 2021). The detection of *Bsal* in the heated Konin lakes during winter suggests that warm-winter conditions may provide a favourable environment for this parasite. These findings underscore the need for additional research, particularly regarding host demographics (Bielby et al., 2021), to assess the potential presence and implications of *Bsal* in the heated environments of the Konin lakes.

### Warming-induced shifts in parasite distribution and elevation in richness are mirrored by similar shifts in host taxa

A total of 39 parasite ASVs exhibited a strong preference for specific lake types, indicating distinct parasite assemblages in heated versus non-heated habitats. Of these, 34 were more prevalent in heated lakes, including *Aphelidium desmodesmi* and four other ASVs belonging to Aphelidiaceae. Although the thermal preference of aphelid parasites remains underexplored, *A. desmodesmi* infections in the green alga *Scenedesmus* were previously shown to increase under elevated summer temperatures (22-25 °C) and phosphorus-limiting conditions in a mesocosm study (Alam et al., 2024). Consistent with this, green algal and diatom hosts of *A. desmodesmi*, including *Desmodesmus armatus* (=*Scenedesmus armatus*, ASV 27), have shown a preference for, and higher relative abundance in heated lakes (Beng et al., 2023). *D. armatus* typically shows optimum growth at high temperatures (∼27 °C) (S. Wang et al., 2019), matching the summer water temperatures of the heated Konin lakes. This suggests that warming may increase aphelid parasite richness both by directly facilitating epidemics at elevated temperatures and indirectly by promoting host species growth. Similarly, the parasitic dinoflagellate *Dissodinium pseudolunula* (ASV 110), which infects copepod eggs, also showed a preference for heated lakes, suggesting that warming may favor this parasite as well. Conversely, two unidentified chytrid ASVs (34 and 263; Order: Rhizophydiales) showed a preference for non-heated lakes and were associated with the diatom host *Ulnaria* (=*Synedra*), which also showed a preference for control lakes (Beng et al., 2023). The thermal preferences of other indicator parasites, have not been extensively investigated in semi-natural or laboratory settings. Overall, these findings suggest that long-term warming has led to the development of distinct parasite assemblages, likely influenced by host distribution patterns, despite the potential flow of water between heated and non-heated lakes (Zdanowski et al., 2020).

The richness of chytrid parasites in our study (which was elevated in heated lakes) showed a general positive correlation with host richness, despite known variation in the extent of their host-specificity. The host-diversity-begets-parasite-diversity hypothesis is proposed to be driven by host-specificity of parasites (Krasnov & Poulin, 2015). While some chytrid parasites are highly host-specific (Van den Wyngaert et al., 2018), other evidence suggests that the specificity might be flexible, particularly during the initial “attraction” phase when motile zoospores find and attach to potential host cells. The chemical cues that attract motile zoospores, such as dimethylsulfoniopropionate (DMSP), are derived from algal cells and are non-specific, suggesting that the attraction might not be limited to particular algal species (Frenken et al., 2017; Vallet, 2024). Additionally, genotype-specificity of some chytrids may shift across thermal regimes, as seen in *Zygorhizidium planktonicum* infecting the diatom *A. formosa* (Gsell et al., 2013). Nevertheless, the positive correlation between host and parasite richness in our study suggests some degree of specificity between chytrid parasites and their compatible host species (or genotypes). Increased host richness may also support a greater richness of generalist parasites. Chytrids that exhibit a ‘generalist’ versus ‘specialist’ infection strategy often occur at different times of the year (Kagami et al., 2021; Van den Wyngaert et al., 2022). For example, the seasonal abundance of chytrid parasites often corresponds with fluctuations in their specific phytoplankton host taxa (e.g. diatoms, cyanobacteria) (Rasconi et al., 2012), suggesting that parasites may “track” the most abundant host species. During spring diatom blooms, chytrids tend to specialize on the dominant diatom species, while an even host community towards the end of the bloom supports the persistence of generalist chytrids (Sassenhagen et al., 2023). Thus, for chytrid parasites, increased host richness may support higher parasite richness regardless of whether the parasites are specialist or generalist, potentially at different times of the year, contributing to their overall increased richness in heated lakes.

### Warming alters seasonal richness patterns of parasitic chytrids, but not Peronosporomycetes (=Oomycetes) or Cryptomycota

The richness of chytrid parasites in heated lakes exceeded expectations based on host richness alone during autumn. One potential explanation for this finding is the significantly higher soluble phosphorus content observed in heated lakes during this season (Beng et al., 2023). Phosphate levels can affect both hosts and parasites. Elevated phosphorus is linked to increased infection prevalence of some chytrids, such as *Rhizosiphon crassum* infecting its cyanobacterial host *Anabaena flosaquae* (Rasconi et al., 2012). Conversely, low phosphorus levels can reduce the growth rate of diatom hosts such as *A. formosa* without impacting the growth rate of their *Rhizophydium* parasites, leading to epidemics at lower host densities (Bruning, 1991). Heated lakes harbor cyanobacterial blooms in the autumn, which could grow to high densities due to elevated phosphorus, and serve as alternative hosts – albeit none of the chytrid parasites in our study associated with cyanobacterial hosts (see Fig. 1B). Thus, the increase in phosphorus levels in heated lakes during autumn (Beng et al., 2023) could underlie the unexpectedly high chytrid parasite richness in autumn.

Warming is hypothesized to induce shifts in the host range of parasites (Wells & Clark, 2019), or favor generalist parasites due to their ability to infect multiple hosts (Cizauskas et al., 2017). Four indicator parasites - the mussel *Elliptio complanata,* fungus *Ustilago esculenta,* and chytrids *Dinomyces arenysensis,* and *Dangeardia mamillata*, all known host generalists – were more prevalent in heated lakes (see Fig. 2). *E. complanata* (ASV 462) is an ectoparasite (Gendron et al., 2019) that can attach to at least 40 fish host species spanning 14 families (Lellis et al., 2013). *U. esculenta* belongs to a hybridizing species complex of multicellular fungi that parasitize multiple hosts within the *Poaceae* grass family (reviewed in McTaggart et al., 2012). The chytrid *Di. arenysensis* infects at least four genera of dinoflagellates (Lepelletier et al., 2014), whereas *Da. mamillata* infects green algae from at least three genera (Van den Wyngaert et al., 2018). Other indicator taxa, including the aphelid *Aphelidium desmodesmi* and the chytrids *Quaeritorhiza haematococci* and *Rhizophydium chlorogonii* (=*Aquamyces chlorogonii*), were also more prevalent in heated lakes, but their host ranges remain unknown.(Beng et al., 2023)(Hutorowicz, 2006)(Dunn et al., 2008)Further research on host ranges is needed to test (Agosta et al., 2010; Wells & Clark, 2019) the potential for warming to favour generalist parasites (Cizauskas et al., 2017).

In contrast with chytrids, warming did not alter seasonal richness patterns of oomycetes or Cryptomycota. Oomycete richness peaked during spring and summer in the current study, potentially aligning with the elevated abundance of airborne (Lang-Yona et al., 2018) and freshwater (Hallett & Dick, 1981) oomycetes documented in spring and summer. However, there are no studies investigating seasonal variation in oomycete parasite species richness, especially among freshwater taxa. Cryptomycota are known to hyperparasitize chytrids (reviewed in Bermúdez-Cova et al., 2023), and are thus proposed to impact phytoplankton communities (Gleason et al., 2017). In the current study, Cryptomycota richness peaked in autumn and winter. This could potentially be linked with the peak in parasitic chytrid richness in autumn; however, further investigation at greater temporal resolution is required to confirm this. Furthermore, not all Cryptomycota hyperparasitize chytrids: some are also reported to infect amoebae (Seto et al., 2023), highlighting how research on the host ranges or thermal preferences of these enigmatic groups (Cryptomycota and oomycetes) is currently insufficient to model expectations of their potential responses to warming.

### Limitations of eDNA-based monitoring for disease ecology

eDNA metabarcoding is a powerful tool for biodiversity-monitoring, but has well-documented limitations (Garlapati et al., 2019). Two key limitations are the lack of quantitative estimates of taxon abundance and the inability to distinguish between the presence of live organisms and that of dormant life stages such as spores. Overcoming these issues by complementing eDNA metabarcoding with quantitative and organismal approaches could improve our insight into the effects of warming on disease.

One important aspect of disease ecology is the abundance of hosts, which cannot be estimated from a metabarcoding approach alone, and which significantly impacts parasitism in the environment. In our study, heated lakes exhibited stronger seasonal variation in parasite richness than control lakes, which may be due to the unexpectedly high chytrid parasite richness in autumn. This suggests that seasonal ecosystem-wide processes driven by warming may contribute to the peak in parasite richness in autumn. Moreover, the higher parasite richness and distinct species assemblages in heated lakes, combined with previously reported elevation in host richness in these environments (Beng et al., 2023), raises the possibility that warming may foster novel host-parasite encounters. For instance, heated lakes harbor the invasive tape grass *Vallisneria spiralis*, which peaks in biomass during autumn (Hutorowicz, 2006). *Vallisneria* macrophytes can increase the abundance of their epibiont taxa that include green algae, diatoms, and dinoflagellates (Dunn et al., 2008), all of which serve as phytoplankton hosts for a broad range of chytrid parasites (see Fig. 1B). We speculate that this may promote abundance of hosts and generate ‘hotspots’ for novel encounters between potential host species and motile zoospores of generalist parasites. This conceivably increases the likelihood of host shifts, and potentially contributes to the high chytrid parasite richness in heated lakes observed in autumn. Such hypothesized outcomes of warming (Agosta et al., 2010) require quantifying host and parasite abundance in addition to richness to gain a complete understanding of the effects of warming on disease.

Another limitation of eDNA is that detected taxa may not be biologically active, or may originate from dormant dispersal stages. This aligns with our observation that 27% of parasite ASVs from the protist dataset and 41% from the fungal dataset were unexpectedly associated with terrestrial hosts, including plants, animals and humans (See Fig. 1). As one example, the fungal ASV 259 shared 100% sequence similarity with *Botrytis cineria* (Phylum: Ascomycota, GenBank AY544651), a pathogen that disperses through airborne spores and causes grey mold in terrestrial plant hosts (Williamson et al., 2007). *B. cineria* has also been shown to cause severe developmental abnormalities and fitness loss in aquatic organisms like the zebrafish *Danio rerio* (Shi et al., 2023, see Table S4). This illustrates how some fungal parasites, although typically associated with terrestrial hosts, may impact aquatic species. Despite such a possibility, we cannot rule out that this taxon was detected from dormant spores. eDNA-based detection must therefore be complemented with organismal approaches such as microscopic counts to confirm the presence and abundance of live parasites *in situ*.

## Conclusion

Our study revealed a vast diversity of eukaryotic parasitic plankton species, with warming shown to increase parasite richness for some groups such as chytrids and aphelids. Parasites exhibited distinct habitat preferences, favouring either heated or non-heated lakes, resulting in unique parasite species assemblages. These preferences might be linked to host species distributions. For chytrid parasites, increased richness in heated lakes can be partially attributed to greater host richness in these environments. For chytrid parasites, warming amplified seasonal variation in richness and increased the strength of correlation between host and parasite richness. Habitat preferences of parasites suggest that warming can shape species assemblages by impacting both abiotic factors (such as thermal conditions in heated lakes) and biotic factors (such as host richness), demonstrating how parasite richness and assemblages may shift under global change.

## Supporting information

supplementary R scripts

## Declarations

### Ethics approval and consent to participate

Not applicable

### Consent for publication

Not applicable

### Availability of data and material

Raw sequence data for the first sequencing run are available in the European Nucleotide Database (ENA) under BioProject ID PRJEB54709 and those of the second run are available at NCBI Sequence Read Archive (SRA) under BioProject ID PRJNA1201567. Reproducible R code for bioinformatic processing is provided in supplementary R scripts.

### Competing interests

The authors declare that they have no competing interests.

### Funding

This work was funded by a joint Beethoven Life-1 grant from the German Science Foundation (WO 1587/9-1 to JW) and the National Science Centre, Poland (2018/31/F/NZ8/01986 to SC).

### Authors’ contributions

JW and SC secured funding and designed the project with MTM. JW and SC collected eDNA samples. AR and KCB performed bioinformatic processing of samples. AR performed ecological analyses and wrote the manuscript with input from all co-authors.

## Acknowledgement

We thank Elisabeth Funke for conducting metabarcoding library preparation, and Sarah Sparmann and Susan Mbedi at the Berlin Center for Genomics in Biodiversity Research (BeGenDiv: https://begendiv.de/) for sequencing support. We further thank the Disease Evolutionary Ecology, and Molecular Ecology & Genomics groups at IGB Berlin for constructive feedback, as well as Nedim Tüzün for stimulating discussions. We are also grateful to two anonymous reviewers whose constructive feedback significantly improved our manuscript.

## Notes

### Competing Interest Statement

The authors have declared no competing interest.

### Summary of Updates

The statistical analyses have been revised to explicitly account for the differences in sequencing depths across the two sequencing runs. The discussion has been rearranged into subsections for clarity.

