## supplementary figures for "Warming increases richness and shapes assemblages of eukaryotic parasitic plankton"

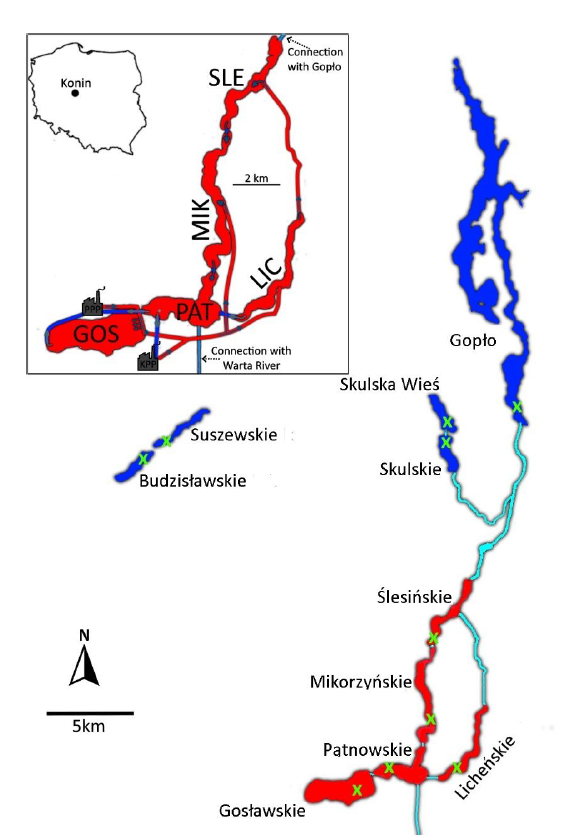
 **Fig. S1** Map indicating study sites with heated lakes indicated in red and control lakes in blue. Green “×” signs denote sampling stations. Water channels connecting the lakes are displayed in cyan. The inset shows a zoom into heated lakes, including the location of the two power plants in Konin and Patnow and the proximity of the lakes to Konin, Poland. The map is reproduced from Dziuba et al 2017.

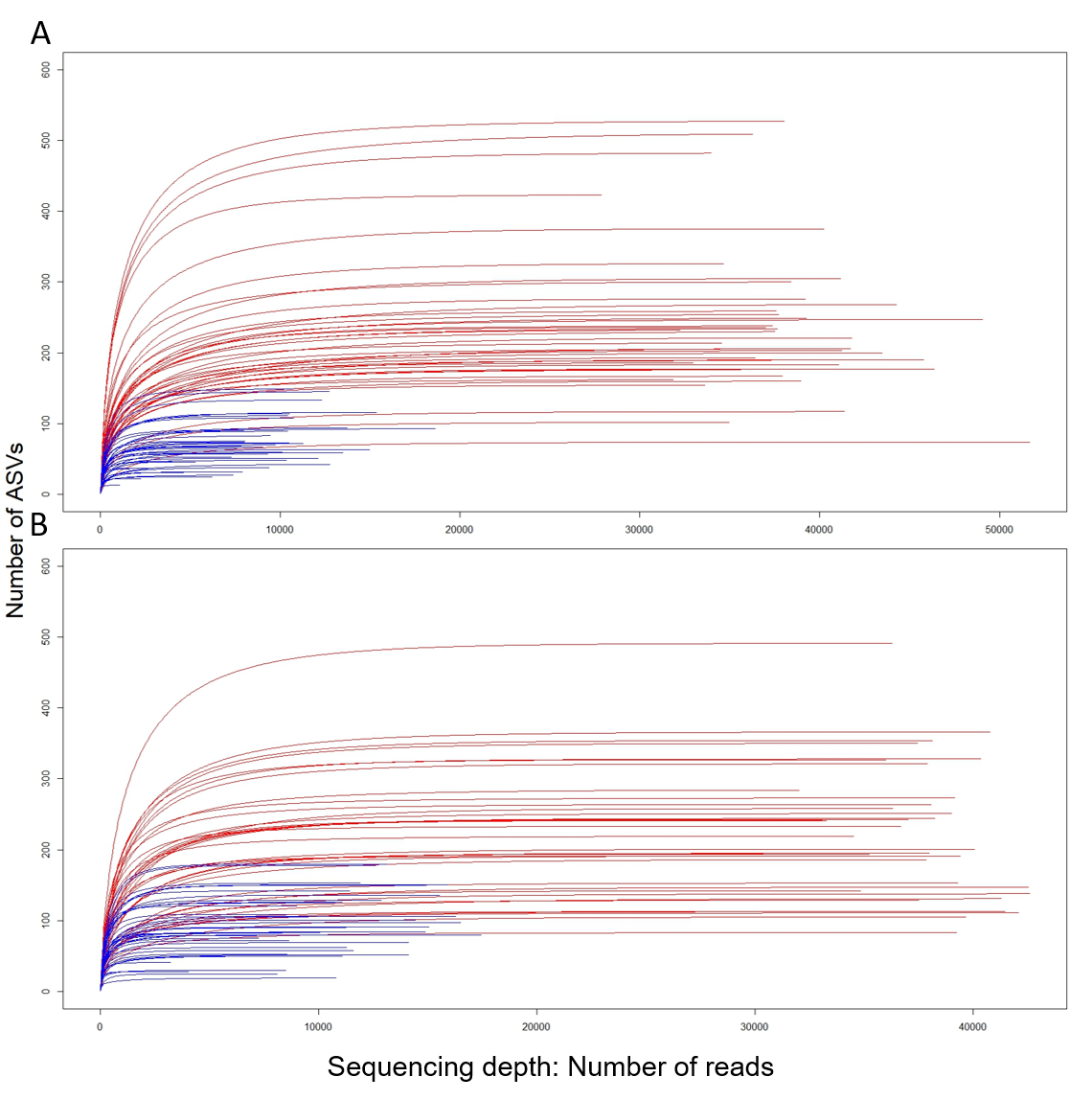

**Fig. S2** Rarefaction curves of samples after pre-processing (see Methods) for (A) the protist dataset and (B) the fungal dataset. Curves are colored-coded by sequencing run (red = run 1, blue = run 2). Fewer reads passed quality control in the second sequencing run; however, all rarefaction curves plateaued, indicating sufficient sequencing depth for both datasets.

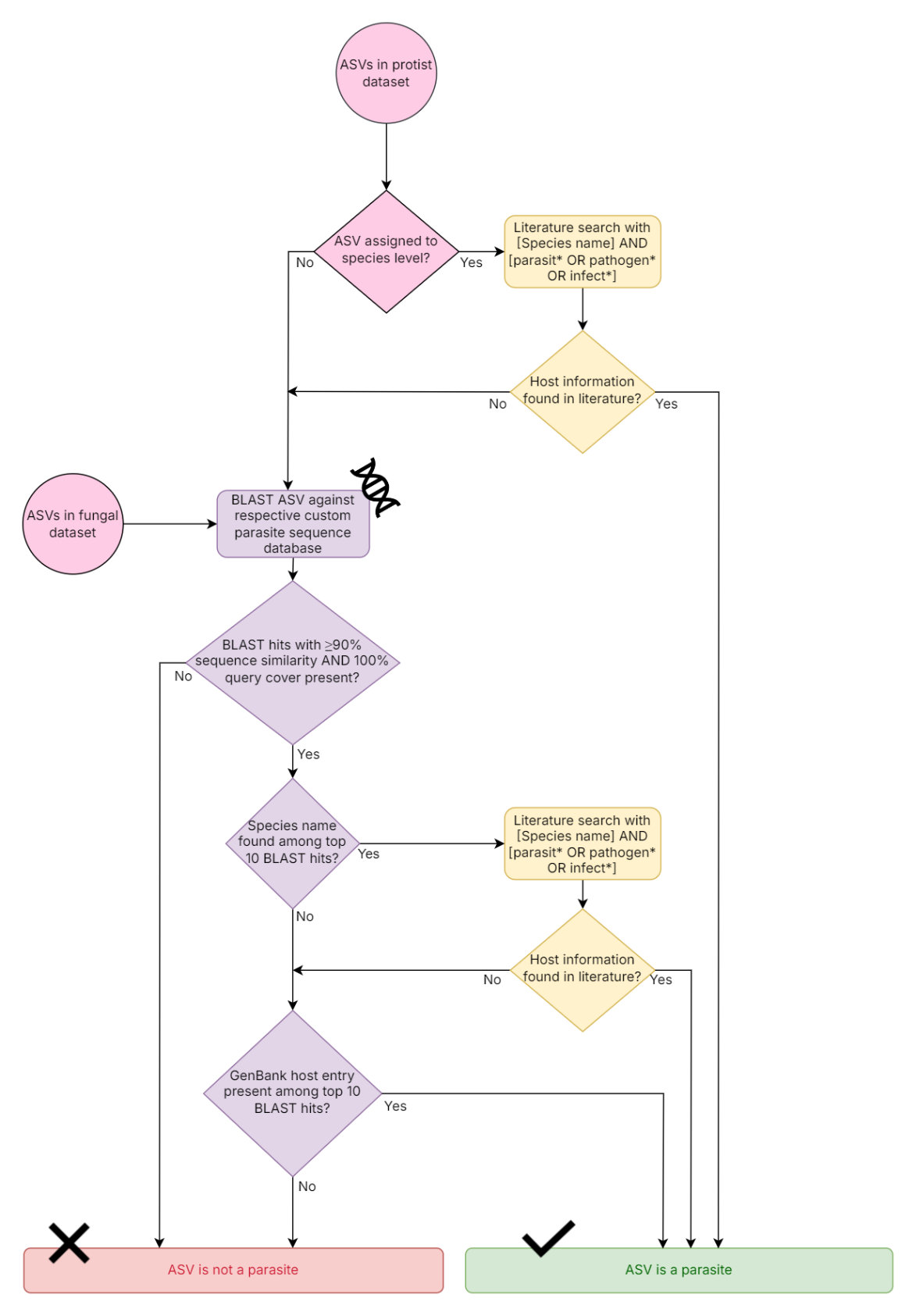

**Fig. S3** Flowchart depicting the classification process of ASVs as parasitic or non-parasitic (see Methods and Beng et al., 2021).Two custom parasite sequence databases were created, containing all NCBI sequences belonging to broad phylogenetic taxa present in the protist (see Fig. S3) and fungal (see Fig. S4) datasets. ASVs from each dataset were BLASTed against their respective custom sequence databases. Following this, GenBank host metadata for parasitic ASVs was reviewed to exclude those with ambiguous host assignments (see Methods and Table S1).

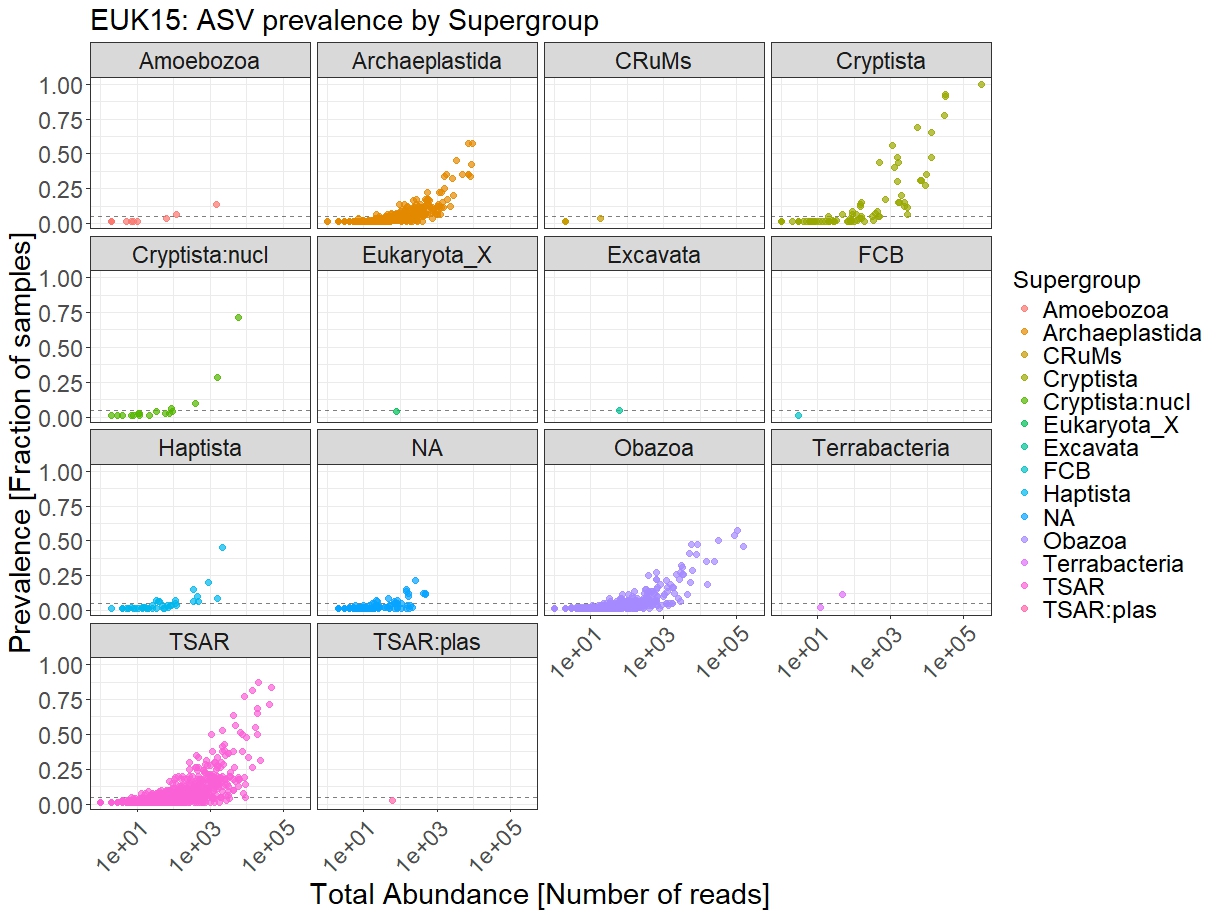

**Fig. S4** Distribution of eukaryotic Supergroups identified in the protist dataset, based on PR2 database assignment. Each point represents an ASV, with the prevalence plotted against read abundance. ASVs are categorized by Supergroup. Sequences belonging to these broad taxa were subsequently downloaded from the NCBI nucleotide database and incorporated into the custom sequence database used in this study (see Methods). A small fraction of reads (0.3% of the total dataset) could not be assigned to any Supergroup and are labelled as “NA”.

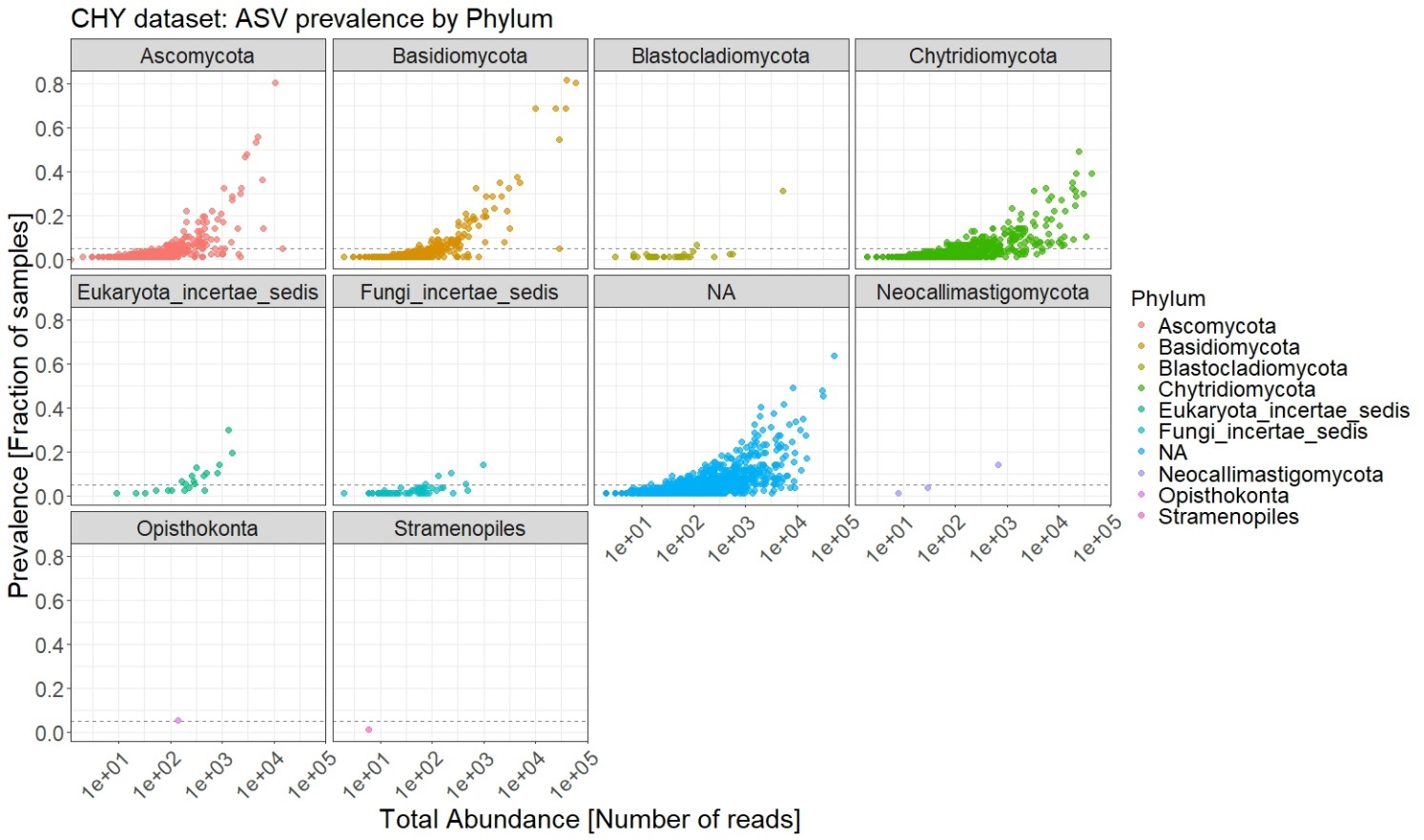

**Fig. S5** All eukaryotic phyla assigned using the RDP LSU database for the fungal dataset. Each point represents an ASV, with prevalence plotted against read abundance. ASVs are grouped by Phylum. A significant portion of the dataset (41.3% of reads, 55% of ASVs) could not be assigned to a specific Phylum and are labelled as “NA”. Thus, a custom NCBI sequence database was developed, to identify chytrids including all above phyla and “Eukaryota_incertae_sedis” and “Fungi_incertae_sedis” (see Methods).

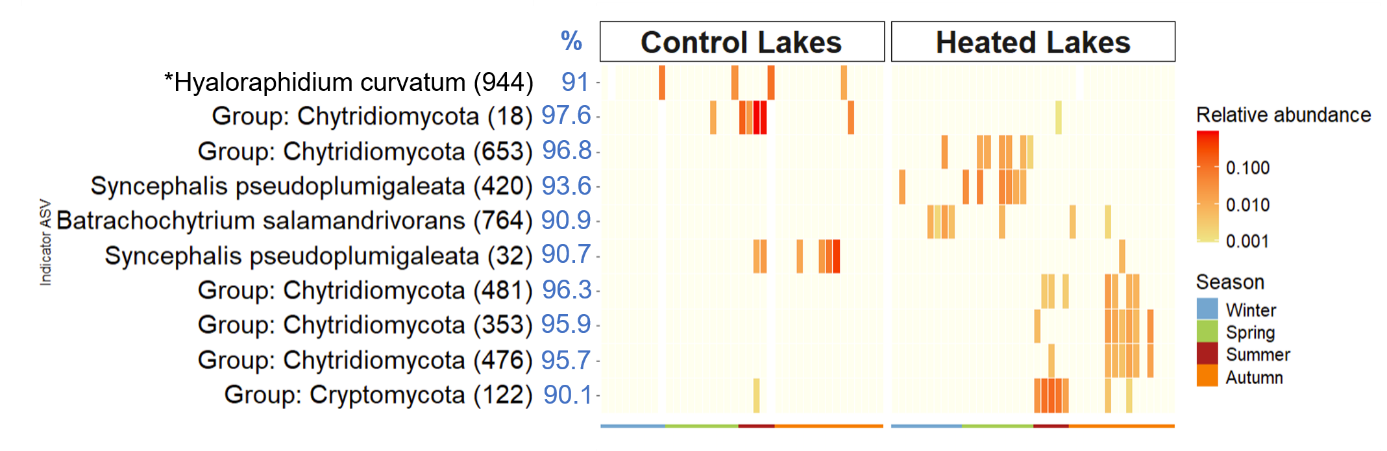

**Figure S6.** Relative abundances of ASVs indicative of control (left panel) and heated (right panel) lakes. Each square represents the relative abundance of indicator ASVs (rows) in individual lakes per year (columns), grouped by the season of sampling indicated by the color of the strip at the bottom. Of the 39 indicator ASVs identified (Table S5), 10 low-prevalence ASVs (present in <20% of samples) are presented here – the remaining 29 are in main text Fig. 2. ASV numbers are specified in brackets. For ASVs in the protist dataset (marked with an asterisk), the lowest taxonomic classification assigned using the PR2 database is reported, with values in the blue “%” column indicating sequence similarity to the BLAST hit used to obtain host information (host entry from GenBank metadata). For ASVs from the fungal dataset, the lowest known classification from the BLAST hit is reported, with the blue “%” values indicating sequence similarity between the ASV and the BLAST hit.

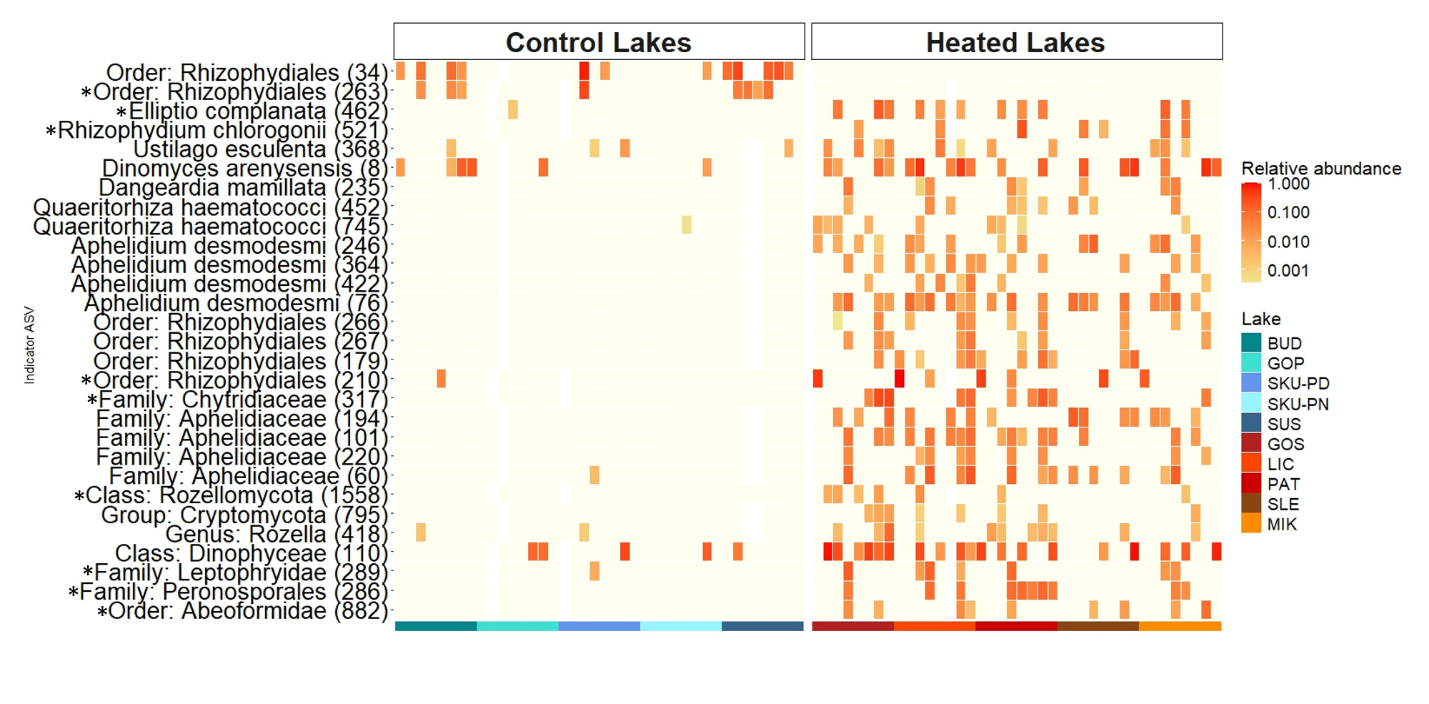

**Figure S7.** Relative abundances of ASVs indicative of control (left panel) and heated (right panel) lakes. This figure is a modification of main text Fig. 2: columns are regrouped to display the relative abundance of indicator ASVs (rows) in individual lakes, which are indicated by the colored strip at the bottom (control lakes BUD = Budzisławskie, GOP = Gopło, SKU-PD = Skulskie, SKU-PN = Skulska Wieś, SUS = Suszewskie, and heated lakes GOS = Gosławskie, LIC = Licheńskie, PAT = Pątnowskie, SLE = Ślesińskie, MIK = Mikorzyńskie). Of the 39 indicator ASVs identified (Table S6), 29 are presented here, as they were present in more than 20% of samples. ASV numbers are specified in brackets. For ASVs in the protist dataset (marked with an asterisk), the lowest taxonomic classification assigned using the PR2 database is reported. For ASVs in the fungal dataset, lowest taxonomic rank of BLAST hit is reported.
