## supplementary R scripts for "Warming increases richness and shapes assemblages of eukaryotic parasitic plankton": EUK15_18S_bioinformatics.R

```
#############################################################################################################################
#Warming increases richness and shapes assemblages of eukaryotic parasitic plankton
#Amruta Rajarajan(*1), Sławek Cerbin(2), Kingsly C. Beng(1, 3§, 4§, 5§), Michael T. Monaghan(1, 6), Justyna Wolinska(1, 6)
# (1) Dept. of Evolutionary and Integrative Ecology, Leibniz Institute of Freshwater Ecology and Inland Fisheries (IGB), Müggelseedamm 301, 12587 Berlin, Germany
# (2) Dept. of General Zoology, Faculty of Biology, Adam Mickiewicz University, Poznań, Poland 
# (3) Focus Group Marine Molecular Ecology, Helmholtz Institute for Functional Marine Biodiversity, University of Oldenburg (HIFMB), Oldenburg Germany
# (4) Alfred Wegener Institute (AWI), Helmholtz Center for Polar and Marine Research, Bremerhaven, Germany
# (5) Thünen Institute of Sea Fisheries, Bremerhaven, Germany
# (6) Institut für Biologie, Freie Universität Berlin, Germany

# *Corresponding author
# §Present address 

# See also Beng et. al (2023) Long-term effects of surface-water temperature increase on plankton communities in artificially heated lakes: insights from eDNA metabarcoding
# https://advance.sagepub.com/doi/full/10.22541/au.168639128.85081228/v1  
#############################################################################################################################

rm(list=ls())
setwd("./Data/Raw/EUK15")

#Load packages
library(Biostrings)
library(Rcpp)
library(dada2)
library(ShortRead)
library(ggplot2)

#set file path
path_EUK15 <- "./Data/Raw/EUK15"  ## Path to the directory containing the fastq files.
list.files(path_EUK15)

#Generate matched lists of the forward and reverse read files
fnFs_EUK15 <- sort(list.files(path_EUK15, pattern = "_R1_001.fastq.gz", full.names = TRUE))
fnRs_EUK15 <- sort(list.files(path_EUK15, pattern = "_R2_001.fastq.gz", full.names = TRUE))

#Identify primers 18S
FWD_EUK15 <- "CCAGCASCYGCGGTAATTCC"
REV_EUK15 <- "ACTTTCGTTCTTGATYRA" 

#Verify the presence and orientation of these primers in the data
allOrients <- function(primer) {
  require(Biostrings)
  dna <- DNAString(primer)
  orients <- c(Forward = dna,
               Complement = Biostrings::complement(dna),
               Reverse = reverse(dna),
               RevComp = reverseComplement(dna))
  return(sapply(orients, toString))
}
FWD.orients_EUK15 <- allOrients(FWD_EUK15)
REV.orients_EUK15 <- allOrients(REV_EUK15)
FWD.orients_EUK15
REV.orients_EUK15

#“pre-filter” the sequences just to remove those with Ns, but perform no other filtering.
fnFs.filtN_EUK15 <- file.path(path_EUK15, "pre-filtN", basename(fnFs_EUK15)) # Put N-filterd files in filtN/ subdirectory
fnRs.filtN_EUK15 <- file.path(path_EUK15, "pre-filtN", basename(fnRs_EUK15))
filterAndTrim(fnFs_EUK15, fnFs.filtN_EUK15, fnRs_EUK15, fnRs.filtN_EUK15, maxN = 0, multithread = TRUE, matchIDs = TRUE)

#Count the number of times the primers appear in the forward and reverse read
primerHits <- function(primer, fn) {
  # Counts number of reads in which the primer is found
  nhits <- vcountPattern(primer, sread(readFastq(fn)), fixed = FALSE)
  return(sum(nhits > 0))
}

rbind(FWD.ForwardReads_EUK15 = sapply(FWD.orients_EUK15, primerHits, fn = fnFs.filtN_EUK15[[1]]), 
      FWD.ReverseReads_EUK15 = sapply(FWD.orients_EUK15, primerHits, fn = fnRs.filtN_EUK15[[1]]), 
      REV.ForwardReads_EUK15 = sapply(REV.orients_EUK15, primerHits, fn = fnFs.filtN_EUK15[[1]]), 
      REV.ReverseReads_EUK15 = sapply(REV.orients_EUK15, primerHits, fn = fnRs.filtN_EUK15[[1]]))

cutadapt <- "./Data/Raw/cutadapt/cutadapt.exe"
system2(cutadapt, args = "--version")

path.cut_EUK15 <- file.path(path_EUK15, "cutadapt")
if(!dir.exists(path.cut_EUK15)) dir.create(path.cut_EUK15)
fnFs.cut_EUK15 <- file.path(path.cut_EUK15, basename(fnFs_EUK15))
fnRs.cut_EUK15 <- file.path(path.cut_EUK15, basename(fnRs_EUK15))

FWD.RC_EUK15 <- dada2:::rc(FWD_EUK15)
REV.RC_EUK15 <- dada2:::rc(REV_EUK15)
# Trim FWD and the reverse-complement of REV off of R1 (forward reads)
R1.flags_EUK15 <- paste("-g", FWD_EUK15, "-a", REV.RC_EUK15) 
# Trim REV and the reverse-complement of FWD off of R2 (reverse reads)
R2.flags_EUK15 <- paste("-G", REV_EUK15, "-A", FWD.RC_EUK15) 

# Run Cutadapt
for(i in seq_along(fnFs_EUK15)) {
  system2(cutadapt, args = c(R1.flags_EUK15, R2.flags_EUK15, "-n", 2, # -n 2 required to remove FWD and REV from reads
                             "-o", fnFs.cut_EUK15[i], "-p", fnRs.cut_EUK15[i], # output files
                             fnFs.filtN_EUK15[i], fnRs.filtN_EUK15[i])) # input files
}

# Check the presence of primers in the cutadapt output
rbind(FWD.ForwardReads_EUK15 = sapply(FWD.orients_EUK15, primerHits, fn = fnFs.cut_EUK15[[1]]), 
      FWD.ReverseReads_EUK15 = sapply(FWD.orients_EUK15, primerHits, fn = fnRs.cut_EUK15[[1]]), 
      REV.ForwardReads_EUK15 = sapply(REV.orients_EUK15, primerHits, fn = fnFs.cut_EUK15[[1]]), 
      REV.ReverseReads_EUK15 = sapply(REV.orients_EUK15, primerHits, fn = fnRs.cut_EUK15[[1]]))

# Run the primer-free sequence files through the DADA2 pipeline
## Processing EUK15_18S sequences using DADA2
## Set working directory make sure all files are present and correctly named

setwd("./Data/Raw/EUK15/cutadapt")  
path_EUK15_cutadapt <- "./Data/Raw/EUK15/cutadapt"
list.files(path_EUK15_cutadapt);

## Extract consistent sample names from forward and reverse fastq files

EUK15_18S.fnFs <- sort(list.files(path_EUK15_cutadapt, pattern="_R1_001.fastq.gz", full.names = TRUE));
EUK15_18S.fnRs <- sort(list.files(path_EUK15_cutadapt, pattern="_R2_001.fastq.gz", full.names = TRUE));
EUK15_18S.sample.names <- sapply(strsplit(basename(EUK15_18S.fnFs), "_"), `[`, 1);
EUK15_18S.sample.names

## visually inspect forward and reverse read quality profiles

plotQualityProfile(EUK15_18S.fnFs[1])
plotQualityProfile(EUK15_18S.fnRs[4])

## Place filtered reads in new subdirectory

EUK15_18S.filtFs <- file.path(path_EUK15_cutadapt, "EUK15_18S.filtered", paste0(EUK15_18S.sample.names, "_F_filt.fq.gz")); 
EUK15_18S.filtRs <- file.path(path_EUK15_cutadapt, "EUK15_18S.filtered", paste0(EUK15_18S.sample.names, "_R_filt.fq.gz"));

## Filter and trim reads based on their quality profiles

EUK15_18S.filter_trim <- filterAndTrim(EUK15_18S.fnFs, EUK15_18S.filtFs, 
                                       EUK15_18S.fnRs, EUK15_18S.filtRs, 
                                       trimLeft=c(0,0), 
                                       truncLen=c(260,200), 
                                       maxN=0, 
                                       maxEE=c(2.6,2.0), 
                                       truncQ=2, 
                                       rm.phix=TRUE, 
                                       compress=TRUE, 
                                       multithread=TRUE);
head(EUK15_18S.filter_trim);

## Learn the specific error-signature of the dataset

EUK15_18S.errF <- learnErrors(EUK15_18S.filtFs, multithread=TRUE);
EUK15_18S.errR <- learnErrors(EUK15_18S.filtRs, multithread=TRUE);

## visually inspect forward and reverse read error models/rates

plotErrors(EUK15_18S.errF, nominalQ=TRUE)
plotErrors(EUK15_18S.errR, nominalQ=TRUE)

## Dereplicate reads and generate new quality-score profiles for each unique sequence based on the average quality scores of each base of all of the sequences that are replicates of it

EUK15_18S.derepFs <- derepFastq(EUK15_18S.filtFs, verbose=TRUE);
EUK15_18S.derepRs <- derepFastq(EUK15_18S.filtRs, verbose=TRUE);

names(EUK15_18S.derepFs) <- EUK15_18S.sample.names;
names(EUK15_18S.derepRs) <- EUK15_18S.sample.names;

## Infer true biological sequences

EUK15_18S.dadaFs <- dada(EUK15_18S.derepFs, err=EUK15_18S.errF, multithread=TRUE);
EUK15_18S.dadaRs <- dada(EUK15_18S.derepRs, err=EUK15_18S.errR, multithread=TRUE);

EUK15_18S.dadaFs[[1]];
EUK15_18S.dadaRs[[1]];

## Merge forward and reverse reads

EUK15_18S.mergers <- mergePairs(EUK15_18S.dadaFs, EUK15_18S.derepFs, EUK15_18S.dadaRs, EUK15_18S.derepRs, verbose=TRUE);
head(EUK15_18S.mergers[[1]]);

## Generate an ASV table

EUK15_18S.run2.seqtab <- makeSequenceTable(EUK15_18S.mergers);
dim(EUK15_18S.run2.seqtab);

table(nchar(getSequences(EUK15_18S.run2.seqtab)));

saveRDS(EUK15_18S.run2.seqtab, "20230721_EUK15_18S.run2.seqtab.rds")

## identify and remove chimera de novo

EUK15_18S.run2.seqtab.nochim <- removeBimeraDenovo(EUK15_18S.run2.seqtab, method="consensus", multithread=TRUE, verbose=TRUE);
dim(EUK15_18S.run2.seqtab.nochim);

table(nchar(getSequences(EUK15_18S.run2.seqtab.nochim)));
sum(EUK15_18S.run2.seqtab.nochim)/sum(EUK15_18S.run2.seqtab);

saveRDS(EUK15_18S.run2.seqtab.nochim, "EUK15_18S.run2.seqtab.nochim.rds");
write.table(t(EUK15_18S.run2.seqtab.nochim), "EUK15_18S.run2.seqtab_nochim.tsv");

## Overview of counts throughout the pipeline

getN <- function(x) sum(getUniques(x));
EUK15_18S.track <- cbind(EUK15_18S.filter_trim, sapply(EUK15_18S.dadaFs, getN), sapply(EUK15_18S.dadaRs, getN), sapply(EUK15_18S.mergers, getN), rowSums(EUK15_18S.run2.seqtab.nochim), round(rowSums(EUK15_18S.run2.seqtab.nochim)/EUK15_18S.filter_trim[,1]*100, 1));
colnames(EUK15_18S.track) <- c("input", "filtered", "denoisedF", "denoisedR", "merged", "nonchim", "final_retained");
rownames(EUK15_18S.track) <- EUK15_18S.sample.names;
head(EUK15_18S.track);

write.csv(EUK15_18S.track, "EUK15_18S.track.csv", quote = TRUE, row.names = TRUE);

#Merge runs 1&2
EUK15_run1 <- readRDS("./EUK15_run1_18S.seqtab.nochim.rds")
EUK15_run2 <- readRDS("./EUK15_18S.run2.seqtab.nochim.rds")
euk15.all <- mergeSequenceTables(EUK15_run1, EUK15_run2)

## Give sequence headers more manageable names (ASV_9, ASV_2...)

EUK15_18S.asv_seqs <- colnames(euk15.all)
EUK15_18S.asv_headers <- vector(dim(euk15.all)[2], mode="character")

for (i in 1:dim(euk15.all)[2]) {
  EUK15_18S.asv_headers[i] <- paste(">ASV", i, sep="_")
}

## Make and write out a fasta of the final ASV sequences
EUK15_18S.asv_fasta <- c(rbind(EUK15_18S.asv_headers, EUK15_18S.asv_seqs))
write(EUK15_18S.asv_fasta, "EUK15_18S.ASVs.fa")
saveRDS(euk15.all, "EUK15_merged.seqtab.nochim.rds")

## Make and write out an ASV table
EUK15_18S.asv_tab <- t(euk15.all)
row.names(EUK15_18S.asv_tab) <- sub(">", "", EUK15_18S.asv_headers)
write.csv(EUK15_18S.asv_tab, "EUK15_18S.ASV_table.csv", quote=F)

# Assign taxonomy
EUK15_18S.seqtab.nochim <- readRDS("EUK15_merged.seqtab.nochim.rds")
EUK15_18S_tax_pr2 <- assignTaxonomy(EUK15_18S.seqtab.nochim, "pr2_version_4.14.0_SSU_dada2.fasta.gz", tryRC=TRUE, multithread=TRUE, minBoot=80, taxLevels = c("Kingdom","Supergroup","Division","Class","Order","Family","Genus","Species"))
saveRDS(EUK15_18S_tax_pr2, "EUK15_18S_tax_pr2.rds")
write.csv(cbind(t(EUK15_18S.seqtab.nochim), EUK15_18S_tax_pr2), "EUK15_18S_tax_pr2_id.tsv", quote=FALSE)
```
